# Social information use across trophic guilds: foraging bumble bees learn from lady beetles

**DOI:** 10.1101/2024.11.07.622477

**Authors:** Marie Muñiz, Brandon T. Meadows, Pablo Lopez, Elinor M. Lichtenberg

## Abstract

Animals use social information gathered by observing other individuals to adjust their behavior to better match the environment and improve fitness. Many insects use social information in various contexts. Bees improve their foraging efficiency by using social information from conspecifics to gauge nectar availability. Bees frequently encounter various heterospecific flower visitors, including those from different trophic groups such as nectaring predators. These heterospecifics may provide valuable information about nectar availability. We determined how bumble bees (*Bombus impatiens*) use visual social information from lady beetles (*Hippodamia convergens*). Lady beetles are predators of small insects, but not bees, and also visit flowers to consume floral nectar to increase their reproductive output. Using laboratory-maintained bumble bees freely flying in arenas, we tested if bees could (1) innately recognize lady beetles as sources of social information about nectar, and (2) learn to use lady beetles to gauge nectar availability. Bees did not innately recognize lady beetles as a source of social information. They correctly learned to associate conspecifics with the presence and absence of food, but learned to associate lady beetles only with the presence of food and not the absence. Our results demonstrate social learning across species and trophic guilds, but suggest limits to when and how bees generalize social information from diverse heterospecific flower visitors.

## INTRODUCTION

Observing others’ interactions with the environment, or using social information, can improve decision making and enhance an individual’s fitness. Adaptive benefits of social information use include predator avoidance (Zamon 2001; Townsend et al. 2012), selective mate choice (Kavaliers et al. 2017; Jones and DuVal 2019), and increased foraging efficiency (Galef and Giraldeau 2001; Chittka and Leadbeater 2005). Social information use can also alter dynamics of populations, ecological communities, and ecosystems (Gil et al. 2018; Riotte-Lambert and Matthiopoulos 2020; Hämäläinen et al. 2023). Most research investigates conspecifics as sources of social information (Buxton et al. 2020). However, animals frequently interact with diverse species (Avarguès-Weber et al. 2013; Goodale and Magrath 2024). For example, multiple bird species cooperatively forage in mixed flocks (Morse 1970) and necrophagous arthropods like beetles, flies, and wasps interact at carrion (Sawyer and Bloch 2020). All of these species may provide social information to one another. Mounting evidence indicates that many animals alter their behavior by learning from heterospecifics (reviewed in Goodale et al. 2010; Gil et al. 2018; Hämäläinen et al. 2023). Understanding how animals use social information from other species is critical. Heterospecific information can directly influence fitness, shape interaction outcomes, and modulate ecological processes. We address this question in the context of bee foraging, asking how bees use social information from heterospecific flower visitors that are in a separate trophic guild.

Foragers can benefit from using social information as it allows them to quickly gain up-to-date information about food resources and thus increase their foraging efficiency. Information includes the location, identity, and quality of novel or familiar resources. Honey bees use their waggle dance to communicate to nestmates the direction and distance of a resource (von Frisch 1967), adult meerkats bring home live prey to teach pups how to identify and handle food (Thornton and McAuliffe 2006), and Norway rats are more likely to sample new food types that did not make conspecifics ill (Galef and Wigmore 1983). Using such information can reduce search effort (Vartparonian and Leu 2024) and food handling times (Danchin et al. 2004).

Animals can adaptively adjust their information use based on environmental conditions and characteristics of the information source. For example, annual killifish rely more on chemical social information when water is too murky to use visual information (Reyes Blengini et al. 2018). Increasing evidence shows that animals can also modulate their behavior depending on the species providing the information. Stingless bees will use species-specific chemistries and concentrations of encountered recruitment pheromones to avoid costly fights over food sources (Nieh et al. 2004; Lichtenberg et al. 2011; Lichtenberg et al. 2014), and birds will gauge the quality of nesting habitat based on heterospecifics’ clutch sizes (Forsman and Seppänen 2011).

Bees are an excellent system for studying plastic use of social information. Multiple bee species – including social bumble (e.g., Leadbeater and Chittka 2007), honey (e.g., Giurfa and Núñez 1992), and stingless bees (e.g., Lichtenberg et al. 2014), as well as solitary bees in several genera (e.g., Yokoi and Fujisaki 2011) – use social information when foraging. This information can come from the visual presence of another bee on a flower or from olfactory cuticular hydrocarbon “footprints” deposited passively by recent flower visitors. In navigating an ephemeral floral food landscape, bees use social information to decide which flower types to sample and rapidly learn which flower species or individual flowers are rewarding (Leadbeater and Florent 2014). Bees can increase their foraging efficiency by feeding from unfamiliar plant species that are being visited, or were recently visited, by conspecifics (Chittka and Leadbeater 2005; Worden and Papaj 2005; Kawaguchi et al. 2006; Dawson and Chittka 2012). Temperate and tropical bees also adjust responses to social information. When visitation drains floral nectar, as in many natural systems, bees often avoid flowers with visual (Stout and Goulson 2001; Kawaguchi et al. 2007) or olfactory (Giurfa and Núñez 1992; Roselino et al. 2016) cues that indicate recent visitation. In contrast, in experiments where social information indicates a constantly-rewarding flower (e.g., *ad libitum* artificial flowers), bees learn to prefer flowers with social information (e.g., Worden and Papaj 2005; Saleh and Chittka 2006; Gloag et al. 2021). The central role of learning in bees’ social information use (Saleh and Chittka 2006; Leadbeater and Chittka 2009) means that they can use information from introduced species (Gloag et al. 2021).

Because social information use by bees impacts floral visitation patterns, it likely plays a role in determining pollination outcomes for both wildflowers and crops. An estimated 90% of all plants and 75% of crops benefit from animal pollination (Potts et al. 2016). While impacts of social information on pollen movement have not been tested, lab experiments indicate potential for such impacts. For example, foraging in the presence of experienced demonstrators may reduce visitation to plant species that do not provide nectar but mimic rewarding plant species (Baude et al. 2008). Modeling suggests that adaptive foraging promotes diverse and robust pollination networks (Valdovinos et al. 2013).

There is a distinct possibility that many different flower visitors, including species in other trophic groups such as predators, provide valuable social information to foraging bees. Bees commonly interact with diverse insects at flowers. Floral nectar and pollen attract many non-bee pollinators such as butterflies and hover flies (Rader et al. 2020; Ollerton 2021; Lichtenberg et al. 2025). Bumble and honey bees reject flowers recently visited by hover flies (Reader et al. 2005). Additionally, many predator and parasitoid insects increase their reproductive success by consuming nectar (Tylianakis et al. 2004; Lundgren 2009) and thus spend time at flowers (Lichtenberg et al. 2023). Despite extensive documentation that insects in multiple taxonomic orders and trophic guilds consume floral nectar and pollen, we know little about whether and how these flower visitors alter bee foraging behavior.

We tested how bumble bees (*Bombus impatiens*) respond to visual social information from predatory lady beetles (*Hippodamia convergens*) via two experiments. Lady beetles consume floral nectar and pollen when insect food is scarce (Hagen 1962; Hodek 1973) or to increase their reproductive output (Lundgren and Seagraves 2011). They can be frequent flower visitors (Aristizábal et al. 2013; Lichtenberg et al. 2023) and thus potentially compete with bees. Because they tend to stay on individual flowers longer than bees do (pers. obs.), lady beetles may be frequent sources of social information for foraging bees in at least some systems or times of year. Lady beetles are not predators of bees, so bees should not exhibit anti-predator behavior in response to the presence of lady beetles. Additionally, our focal species is commercially available and can be maintained in lab colonies, so is experimentally tractable.

To determine whether bees can use nectaring predators as sources of social information about nectar, we asked two questions. (1) Do bees innately recognize nectaring predators as sources of social information? Given bees’ failure to innately recognize inorganic cues (Dawson and Chittka 2012) and the strong visual differences between bees and lady beetles, we predicted that beetle-naïve bees would not be attracted to flowers occupied by lady beetles. Answering this question is important for interpreting bee behavior when given the opportunity to learn from lady beetles. (2) Can bees learn to use nectaring predators as indicators of nectar availability? Given bees’ strong associative learning abilities (Saleh and Chittka 2006; Leadbeater and Chittka 2009) and documentation of bees learning from other pollinators (Reader et al. 2005), we predicted that bumble bees would learn to use the presence or absence of lady beetles to gauge nectar availability.

## METHODS

### Study insects

We obtained commercial bumble bee colonies from Biobest (USA) and housed them in a plastic nest box (21×27×12.5 cm) connected by plastic tubing to a foraging arena with a wood frame, mesh walls, and a floor painted green (91×63×60 cm; Glidden™ essentials® low odor flat “Succulent Leaves” interior paint; Fig. 1A). We divided the arena into two sides with a solid, opaque wall. One side was used for daily foraging activities and the other side for experiments. We fed bees *ad libitum* 30% (g sugar/g solution) sucrose solution in the foraging arena for 9 hours on weekdays and continuously on weekends, and pollen in the nest box as needed. 30% sucrose is a common maintenance diet for lab-housed bumble bees (e.g., Baude et al. 2011; Avarguès-Weber et al. 2018). We kept bees on a 12-hour photoperiod (light 7:00-19:00) and further illuminated the foraging arena with full spectrum LED lights (Green Light Depot, 40W, 5000K). We maintained the lab at 25°C (as in Francis et al. 2016) and ∼30% relative humidity (as in Mundy-Heisz et al. 2022).

**Figure 1.**
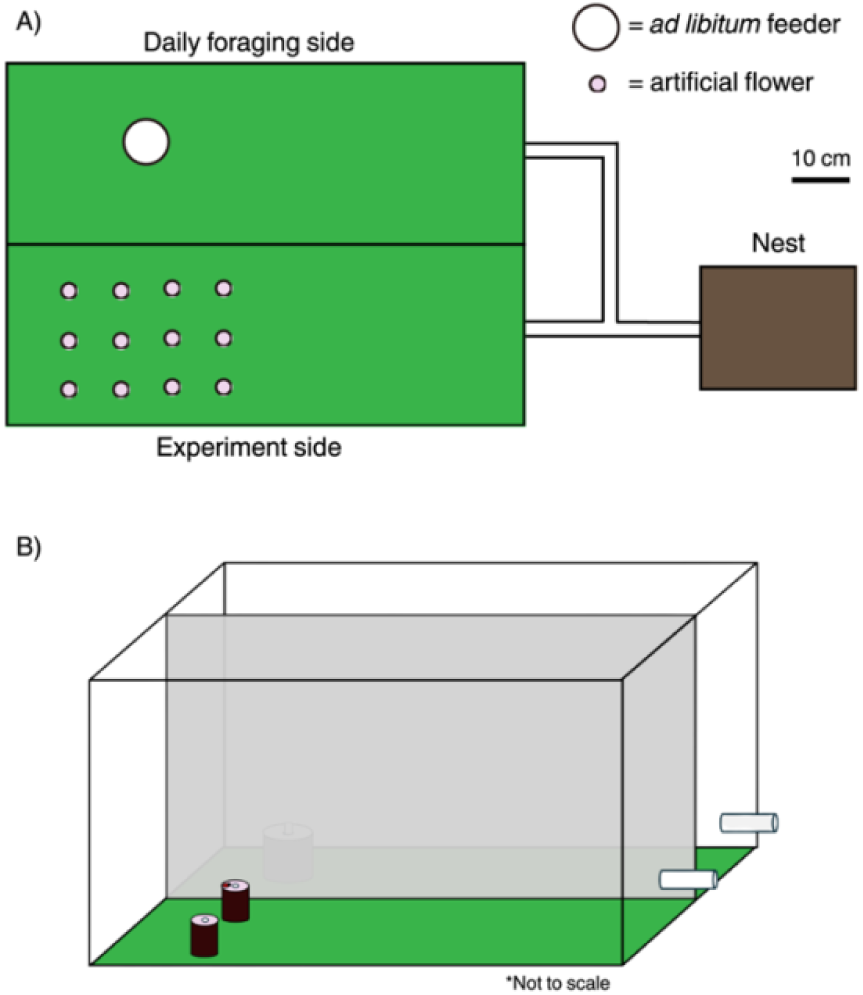
(A) To-scale schematic of a bee nest box connected to a flight arena that is divided into daily foraging and experiment sides. The schematic also shows the bees’ *ad libitum* feeder (white circle) and an array of artificial flowers for the learning experiment (purple circles). (B) Not-to-scale 3D schematic of the flight arena depicting the *ad libitum* feeder on the daily foraging side and an array of artificial flowers for the innate recognition experiment. The wall splitting the two sides was opaque.

We used two different insect species as demonstrators: *H. convergens* lady beetles (purchased live, Clark & Co Organic) and the same bumble bee species used as trial subjects (*B. impatiens*). We presented demonstrators on artificial flowers as pinned dead insects. Pinned dead bees, and even clay shaped and painted to look like bees, elicit similar responses as live demonstrators (Worden and Papaj 2005; Leadbeater and Chittka 2007) and allow precise control of information presented. Demonstrators were ethanol-killed, rinsed and dried with hexane five times to eliminate odors, and pinned. On the fifth hexane wash, bees were transferred to a mesh strainer containing small Kimwipe® balls and gently tumble-dried with a blow dryer to fluff their hair (setae). We cut off pin heads to minimize altering demonstrators’ natural appearance.

### Experimental setup

We constructed artificial flowers from 30mL brown Nalgene bottles with a hole drilled in the lid and a laminated paper disk placed on the top surface (Fig. 1B). Through this hole, we inserted either a cotton wick for training prior to experimentation or a shallow well during experiments. The cotton wick allowed bees to forage *ad libitum*. The shallow well held only a small amount of sucrose solution and could be emptied by a bee in a single visit, simulating drained flowers.

Prior to experiment trials, we familiarized bees with the artificial flowers by letting bees forage on an array of six artificial flowers containing *ad libitum* 30% sucrose solution, each bearing a laminated disc of white printer paper. To standardize prior social information exposure, training flowers contained a pinned bumble bee demonstrator. All bees learned to associate nestmates with sucrose solution during daily feeding; pinned bumble bees on training flowers prevented them from learning to associate empty flowers with a demonstrator. We uniquely identified actively foraging bees with queen number tags (Betterbee®) attached with super glue to their thorax. Bees that visited the artificial flower array at least three times were eligible for subsequent trials.

Two experiments presented bees with artificial flowers containing “orchid” purple discs (Wausau Papers Exact® Multipurpose printer paper) and bumble bee or lady beetle demonstrators. In the innate recognition experiment, flowers contained 150µL of tap water. In the learning experiment, flowers contained 15µL of 50% sucrose solution or tap water. Flowers were washed with residue-free lab detergent and air dried prior to use in experiments, to remove olfactory cues. During a trial, a single bee foraged alone on the “experiment” side of the flight arena. We controlled access to the arena using a series of plastic gates along the tubing that connected the nest box with the flight arena. We defined a flower visit as a bee landing on and extending her proboscis into the well of an artificial flower. A bee that completed a trial was not re-tested. We terminated a trial and manually returned a forager to the nest if she did not interact with flowers within 10 min of entering the flight arena. Pinned demonstrators were not used again for at least five days, to allow dissipation of any footprints that might accumulate from a foraging bee landing on a demonstrator.

### Innate recognition of nectaring predators as sources of social information

In this experiment, bees chose between two artificial flowers with either a bumble bee (n=72 bees) or lady beetle (n=70) demonstrator present on one of the flowers (bees from 11 colonies). Flowers were placed 3 cm apart on an axis perpendicular to the flight arena entry tube, and their position was randomized. A trial began when a bee entered the arena and ended as soon as she visited her first flower. Fifteen bees that only landed on top of flowers but did not extend their proboscis into the well were excluded from analysis and the sample sizes given above.

To evaluate whether bees innately recognize nectaring predators as sources of social information about nectar, we compared whether each bee first visited the occupied flower to chance expectations of visiting an occupied flower (0.5) using two-tailed binomial tests. We separately analyzed trials with bumble bee and lady beetle demonstrators. All analyses were conducted in R v4.3.1 (R Core Team 2023).

### Learning nectar availability from nectaring predators

In this experiment, we first provided foragers the opportunity to learn to associate a demonstrator species with nectar presence or absence, then tested whether the bee learned the association (7 colonies, 3-15 bees/colony). A bee underwent one of four treatments during this experiment: (1) bumble bee demonstrators indicate nectar present (n=11 bees), (2) lady beetle demonstrators indicate nectar present (n=11), (3) bumble bee demonstrators indicate nectar absent (n=11), (4) lady beetle demonstrators indicate nectar absent (n=10; Fig. 2). During a “learning phase”, we presented the bee with an array of six rewarding flowers each containing 15µL of 50% sucrose solution and six non-rewarding flowers each containing 15µL of tap water. Flowers were haphazardly placed in a 4 x 3 grid (Fig. 1A). Bees foraged on this array for three bouts (as in Dawson and Chittka 2012; Hemingway and Muth 2022), with the flower array haphazardly rearranged among bouts (Fig. 2). Bees could visit the same flower within a bout, so flowers were promptly refilled after a visit to ensure bees were always exposed to the same learning conditions. A bout ended when the bee was done foraging and returned to the nest. All bees explored both rewarding and unrewarding flowers; in the first bout, they visited rewarding flowers an average 9 times and unrewarding flowers 8 times. After a bee successfully completed three learning bouts, we conducted a test bout where all 12 flowers contained water. We recorded the number and order of visits to flowers with and without demonstrators during this “test phase” (Fig. 2). Bees that completed three learning bouts followed by a test bout were not tested again.

**Figure 2.**
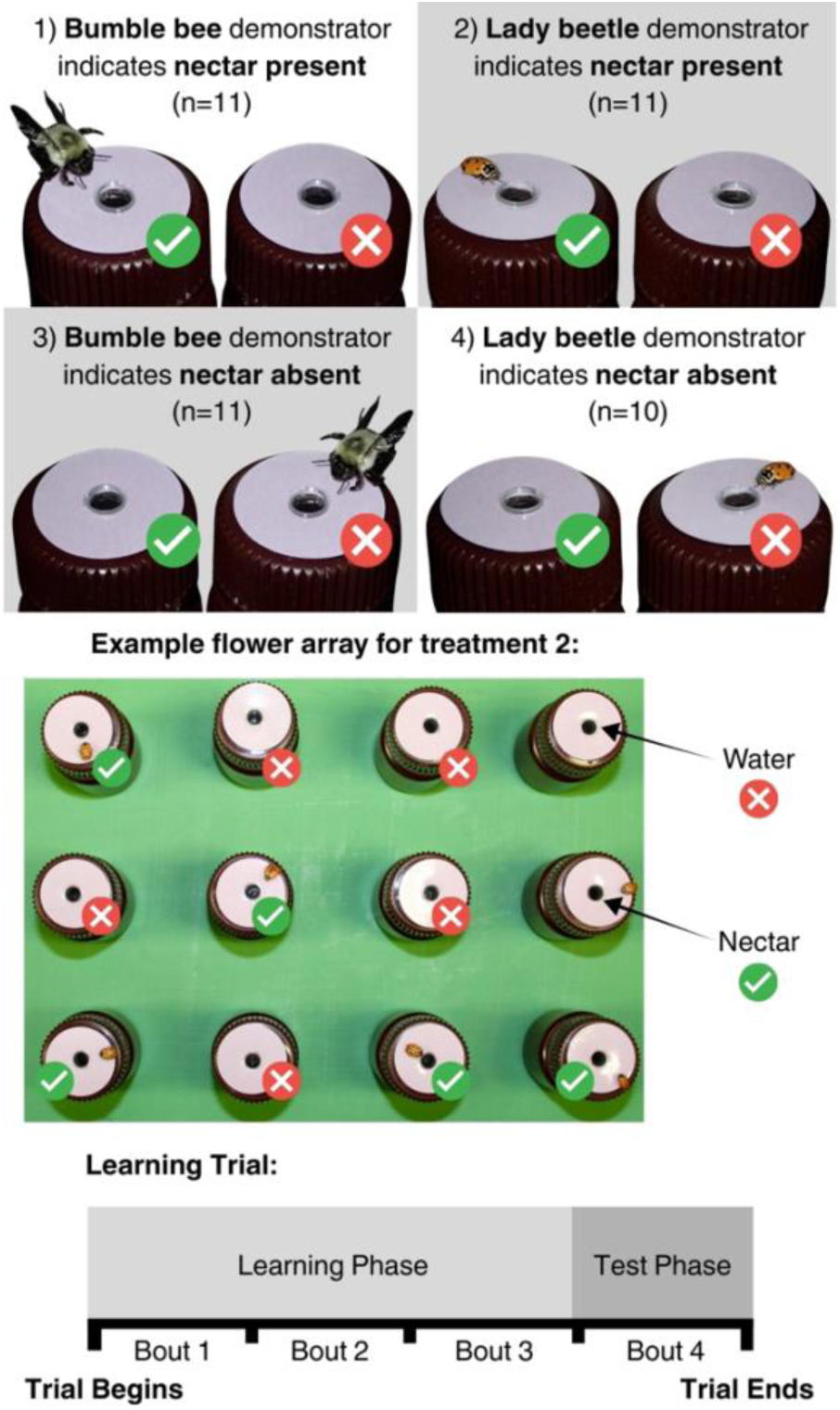
Schematic depicting the learning experiment’s four treatments, an example flower array for treatment 2, and the timeline of a learning trial. Green checkmarks signify when sucrose solution was present in the artificial flowers and red strikes signify when water was present in the flowers. Treatments were only deployed during the learning phase. In the test phase, all flowers contained water.

To evaluate whether bees learned to use nectaring predators to gauge nectar availability, we assessed learning based on test phase behavior in two ways. (1) We determined whether bees selected a flower with a demonstrator on their first flower visit. To determine if treatment affected this behavior, we used a logistic regression with flower type first visited as the response variable; demonstrator species, whether the demonstrator indicated rewarding or non-rewarding flowers, and their interaction as fixed effects; and colony as a random effect (lme4 package, Bates et al. 2015). We also compared a bee’s first flower visit to chance expectation (0.5) of visiting the flower type that was previously rewarding using two-tailed binomial tests, with a separate test for each treatment. (2) We asked whether bees preferentially visited flowers that previously indicated a reward during the first 10 visits in the test phase. To determine if treatment affected this behavior, we used a logistic regression with proportion of visits to previously-rewarding flowers as the response variable; demonstrator species, whether the demonstrator indicated rewarding or non-rewarding flowers, and their interaction as fixed effects; colony as a random effect; and a weight of 10 visits for each proportion. We used two-tailed *t*-tests to determine if the proportion of a bee’s first 10 visits to previously-rewarding flowers differed from chance expectation (0.5). For *t*-tests, we transformed proportions using Anscombe’s arcsine transformation (Zar 1999) to meet parametric assumptions. All regressions met model assumptions.

## RESULTS

### Innate recognition of nectaring predators as sources of social information

As predicted, bees did not innately recognize nectaring predators as a source of social information about food (Fig. 3). Bumble bees preferred flowers occupied by a (familiar) conspecific demonstrator (binomial test: *p*=0.001). They did not show this preference for flowers occupied by (novel) heterospecific lady beetles (binomial test: *p*=0.4).

**Figure 3.**
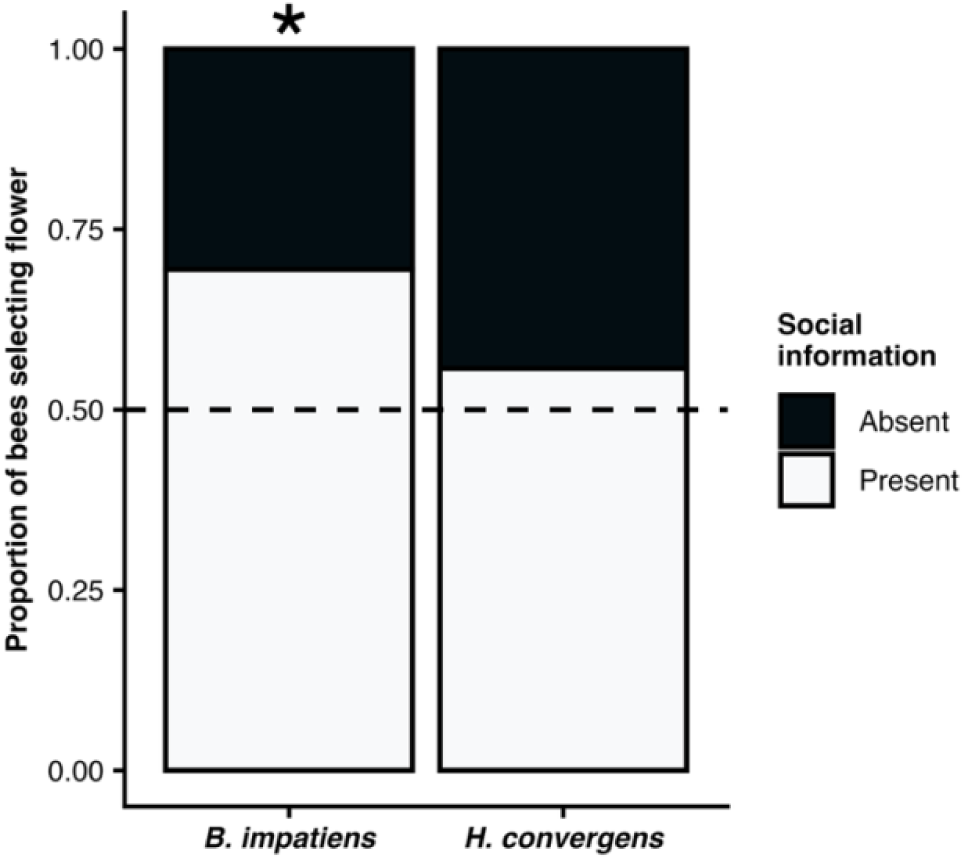
Proportion of bees selecting a flower where social information from a conspecific (*B. impatiens*) or heterospecific (*H. convergens*) demonstrator was present (white) or absent (black). Star indicates a difference from chance (0.5) determined via 1-sample t-tests. Photos © Clay Bolt (*B. impatiens*) and Mike Quinn (*H. convergens*), used with permission.

### Learning nectar availability from nectaring predators

Bees learned that lady beetles could provide social information about nectar availability, but in limited contexts. We first assessed whether bees selected the previously-rewarding flower type (with or without demonstrators) on their first flower visit of the test phase (Fig. 4A). When a conspecific demonstrator was paired with a reward in the learning phase, bumble bees preferred flowers with conspecifics in the test phase (binomial test: *p*=0.01). When a conspecific demonstrator was paired with the absence of a reward, bees did not prefer either flower (binomial test: *p*=1). When a heterospecific demonstrator was paired with either a reward or the absence of a reward, bees did not exhibit evidence of learning in their first flower choice (binomial test: *p*=0.55, 0.34, respectively). Responses to demonstrators on a bee’s first visit differed with demonstrator species, but not with whether the demonstrator previously indicated rewarding or non-rewarding flowers (Table 1).

**Table 1.** Likelihood ratio test results for bees’ (A) first visit and (B) first 10 visits to flowers in the test phase.

| Model Term | $\chi^2$ | df | $p$-value |
| --- | --- | --- | --- |
| **(A) First Visit** |  |  |  |
| Demonstrator Species | 8.25 | 2 | 0.01 |
| Cue Association | 7.76 | 2 | 0.06 |
| Demonstrator Sp. : Cue Assoc. | 2.27 | 1 | 0.13 |
| **(B) First 10 visits** |  |  |  |
| Demonstrator Species | 9.21 | 2 | 0.01 |
| Cue Association | 2.73 | 2 | 0.26 |
| Demonstrator Sp. : Cue Assoc. | 0.2 | 1 | 0.66 |
| Demonstrator species refers to either *B. impatiens* or *H. convergens*. Cue association refers to whether the demonstrator species indicated rewarding or non-rewarding flowers. |  |  |  |

**Figure 4.**
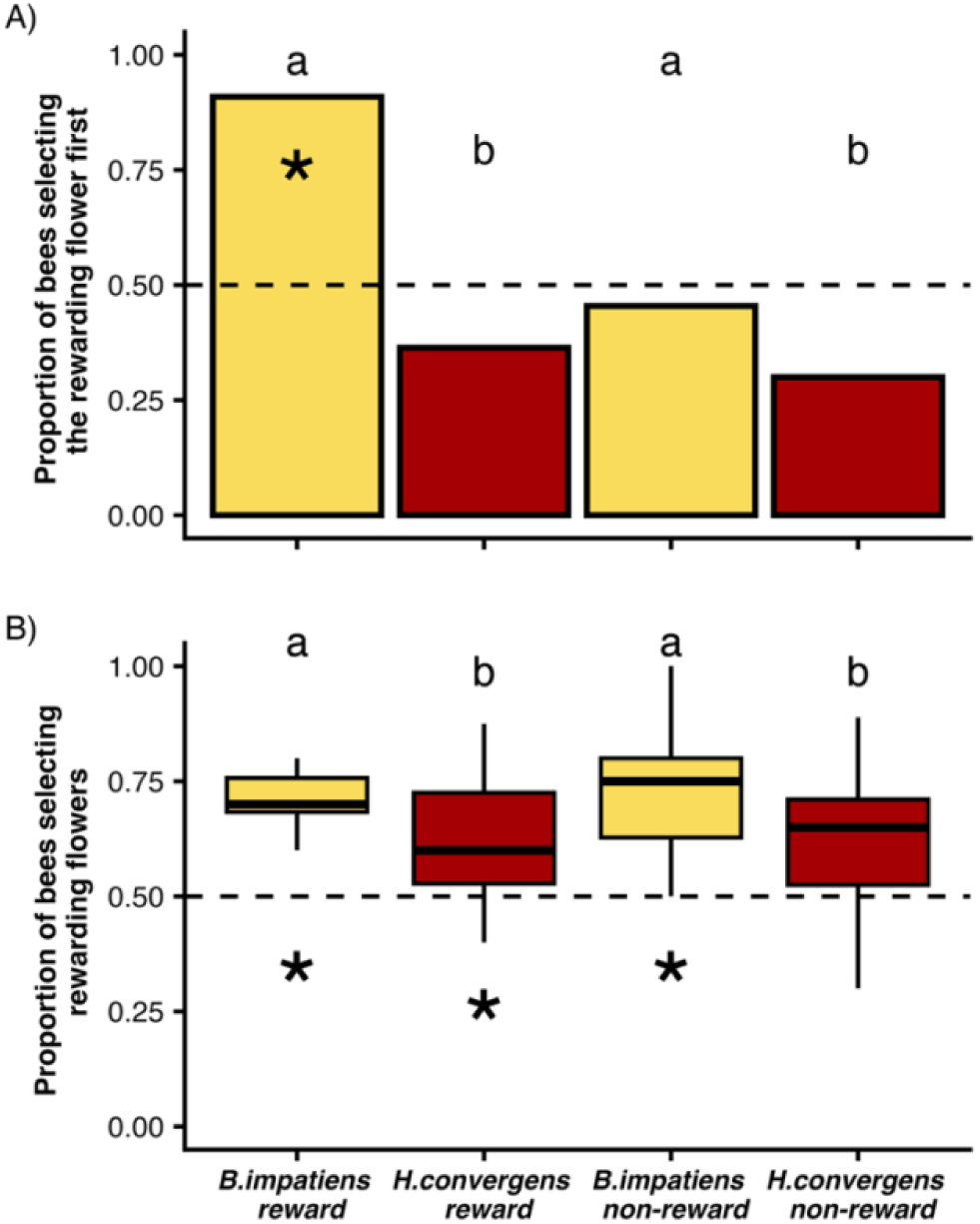
(A) Proportion of bees selecting flowers with social information as their first choice for each respective learning trial type. (B) Proportion of visits to the flowers with social information during a bee’s first 10 choices for each learning trial type. Boxes span the 25th to 75th percentiles, with a wide line at the median, and whiskers extend to the minimum and maximum values. Stars indicate a difference from chance (0.5) determined via 1-sample t-tests. Letters above bars and boxes indicate differences in responses across treatment determined via logistic regression and post-hoc Tukey tests. Bars or boxes with different letters are statistically different from each other.

We also assessed learning using the proportion of a bee’s first 10 visits in the test phase to the flower type that was previously associated with a sucrose reward (Fig. 4B). When a conspecific demonstrator indicated a reward or the absence of a reward in the learning phase, bumble bees preferentially visited the previously-rewarding flower type (*t*_10_=4.70, *p*= <0.0001; *t*_10_=8.22, *p*=<0.0001, respectively). Bees also learned to associate a heterospecific demonstrator with a reward (*t*_10_=3.11, *p*= 0.01). This was not the case when a heterospecific demonstrator previously indicated the absence of a reward (*t*_9_=1.07, *p*= 0.31). In the first 10 visits, bees were more likely to visit rewarding flowers when the demonstrator was a bee. However, whether the demonstrator previously indicated rewarding or non-rewarding flowers did not affect bees’ foraging choices (Table 1).

## DISCUSSION

Social information research has largely focused on information from conspecifics, but animals frequently interact with diverse species and may modulate their behavior in response to such interactions. We investigated how animals use social information from demonstrators in a separate trophic guild. Our results show that bumble bees can learn to use lady beetles as a source of information about floral nectar, in limited contexts. Bees did not innately recognize lady beetles as information sources, in contrast with bee-naïve bees’ preference for flowers occupied by another bee (Kawaguchi et al. 2006). However, foragers did learn to associate lady beetles with the presence, but not the absence, of a sucrose reward. With conspecific demonstrators, bees learned to associate sucrose availability with both the presence and absence of the demonstrator. Thus, bumble bees exhibit flexibility when using social information. Our results suggest that there are limits to how information generalizes across the range of floral visitors. This aligns with research indicating that some solitary bees show no response to other bee species (Yokoi and Fujisaki 2011).

Our innate recognition experiment and learning experiment regressions indicated that bumble bees differentially respond to social information from a conspecific versus a nectaring predator. A flower visitor’s visual appearance may determine whether a bee characterizes the visitor as informative or not, and thus whether there has been selection to recognize or rapidly learn from a given species. Non-informative individuals may be viewed similarly to inanimate objects, which bumble bees have no innate preference for and can be less likely to learn or generalize from (Dawson and Chittka 2012; Avarguès-Weber and Chittka 2014; Romero-González et al. 2020) possibly due to sensory biases (Leadbeater 2015) or contrapreparedness (Dexheimer and Dunlap 2025). Similarly, bumble bees searching for new food sources show no response to woodlice (Romero-González et al. 2020). Animals often ignore or more slowly learn less relevant information (Galef and Whiskin 1997; Albo et al. 2012; Castro and Wasserman 2016). Lady beetles are visually distinct from both bumble and honey bees, with a smaller body size, lack of three articulated body segments, and coloration that is rare for bees. Smaller flower visitors may be harder for bumble bees to see; detecting flowers that are the approximate size of lady beetles (5 or 8 mm) requires greater search time (Spaethe et al. 2001). How a demonstrator’s body size impacts ease of detection and utility as social information are exciting avenues for future research. Lady beetles’ red coloration may also be difficult for bees to distinguish against floral backgrounds since bees lack red-sensitive cones (Menzel et al. 1986; Backhaus and Menzel 1987), although bees do frequently visit red flowers (Sikora et al. 2020).

Alternatively, lady beetles’ bright red and black coloration might warn bees of predation threat. Bumble bees can learn to avoid flowers with cryptic predators such as crab spiders (Ings and Chittka 2009) and flower colors that previously bore a predatory threat (Wang et al. 2013). It is possible that this avoidance extends to bright colors like those displayed by lady beetles. However, our bees showed no innate avoidance of lady beetles. Additionally, in a bee’s environment, bright colors are more often associated with food (flowers) than with predators that are typically cryptic (e.g., spiders) or large (e.g., birds). Thus, a general avoidance by bees of colors typically associated with aposematism (Stevens and Ruxton 2012) seems unlikely.

Another striking result was bees’ failure to respond to lady beetles on non-rewarding flowers despite strong associative learning capabilities (Avarguès-Weber et al. 2011). Simple cognitive phenomena could partially explain this pattern. Prior to experimentation, our bees were exposed daily to social information from other bees at *ad libitum* feeders, equivalent to the *B. impatiens* rewarding treatment. This gave them ample opportunity to learn to associate bumble bees with food. Learning that a novel species (here, lady beetles) indicates nectar-depleted flowers requires generalizing to both a new stimulus and a new functional relationship. Treatments under which bees did learn to use social information altered only the stimulus (lady beetles rewarding) or the function (bees non-rewarding). Learning can be costly (Mery and Kawecki 2004), and generalizing both stimulus and functional relationship may be more difficult to learn than generalizing just one facet of a cue. Similarly, bees show reduced learning performance when switching to flowers that are more difficult to access nectar from (Saleh and Chittka 2006; Sanderson et al. 2006). Bumble bees also exhibit stimulus enhancement when learning only a new functional relationship but local enhancement when learning both a functional relationship and “nonsocial” stimulus (wooden blocks, Avarguès-Weber et al. 2018). Stingless bees may fail to generalize a highly rewarding flower color when the flower pattern changes (Forster et al. 2023).

Competition may also alter behavioral responses to other species (Hämäläinen et al. 2023). Our experimental design used bees whose routine feeding paired bumble bee demonstrators with unlimited reward and low competition. In such an environment, there may be less need to forage efficiently (Norberg 2021) and thus less attention paid to social information. Bees’ first flower choices in the test phase, which did not demonstrate strong preferences, are consistent with this. Wild bees foraging on plants that rapidly replenish nectar (Yokoi and Fujisaki 2009) or contain large amounts of pollen (Stout et al. 1998) also sometimes ignore social information. Bees that switched to a perceived high competition environment in our experiment (bumble bee demonstrators non-rewarding) required multiple visits to show evidence that they learned the association between demonstrator and nectar availability. This is consistent with the hypothesis that bees will avoid flowers where interference competition is likely (Lichtenberg et al. 2010; Baude et al. 2011; Yokoi and Fujisaki 2011) and the observation that, in competitive environments, many animals alter their behavior in response to social information (Hall and Kramer 2008; Périquet et al. 2015; Wüst and Menzel 2017).

Over evolutionary time, there may be selection for recognizing only specific visual characteristics of the demonstrators that a species frequently exploitatively competes with. Certain flower visitors, including very small bees and potentially nectaring predators and parasitoids, consume small enough amounts of nectar that they may not compete with larger bees even when feeding from the same flower. Such flower visitors may be viewed as irrelevant. Inconsistent with this, however, some wild bees and hover flies will reject all occupied flowers independent of demonstrator body size (Yokoi and Fujisaki 2011).

Overall, our results highlight the fascinating and complex nature of social information use across species and trophic guilds. We found that bumble bees are able to learn about flowers’ nectar status by observing a species that is predominantly predatory but also feeds on floral nectar, although the bees do not innately recognize this species as a source of social information. This highlights bees’ high capacity for learning (Chittka and Thomson 2001) and the urgent need to understand how individuals alter their behavior in response to the wide array of other species they encounter in nature (Gil et al. 2018; Lichtenberg et al. 2023; Lichtenberg et al. 2025). Recent research suggests that social information plays a significantly larger role than previously recognized in shaping the evolution of signals, social interactions, and cognition (Avarguès-Weber et al. 2013; Bernal and Page 2022; Kohles et al. 2022), and the ecology of populations, communities, and ecosystems (Gil et al. 2019; Riotte-Lambert and Matthiopoulos 2020; Hämäläinen et al. 2022). For example, one potential outcome of the learning patterns we documented is that bees may use information from lady beetles’ presence on flowers to decide which flower species to visit on a given day, but not for decisions about which individual flowers to visit within that flower species. In this scenario, lady beetle demonstrators may affect a plant species’ pollination rate but not the relative fitness of individual plants within a species. Studying social information use across species and trophic guilds, as in this study, is critical for linking social learning behavior with ecological community dynamics and processes.

## ACKNOWLEDGEMENTS

We thank I. Eastland, A. Gage, A. Kinman, J. Linogao, D. Pedraza, B. Richter, L. Taylor, and M. Vohs for lab assistance; and K. Gotanda, C. Varnon, and three anonymous reviewers for feedback.

